# Structure of the deactive state of mammalian respiratory complex I

**DOI:** 10.1101/165753

**Authors:** James N. Blaza, Kutti R. Vinothkumar, Judy Hirst

## Abstract

Complex I (NADH:ubiquinone oxidoreductase) is central to energy metabolism in mammalian mitochondria. It couples NADH oxidation by ubiquinone to proton transport across the energy-conserving inner membrane, catalyzing respiration and driving ATP synthesis. In the absence of substrates, ‘active’ complex I gradually enters a pronounced resting or ‘deactive’ state. The active-deactive transition occurs during ischemia and is crucial for controlling how respiration recovers upon reperfusion. Here, we set a highly-active preparation of *Bos taurus* complex I into the biochemically-defined deactive state, and used single-particle electron cryomicroscopy to determine its structure to 4.1 Å resolution. The deactive state arises when critical structural elements that form the ubiquinone-binding site become disordered, and we propose reactivation is induced when substrate binding templates their reordering. Our structure both rationalizes biochemical data on the deactive state, and offers new insights into its physiological and cellular roles.

## Introduction

Complex I (NADH:ubiquinone oxidoreductase), a crucial enzyme in oxidative phosphorylation, uses NADH oxidation and ubiquinone reduction to build the proton motive force across the inner mitochondrial membrane, catalyzing respiration and driving ATP synthesis (1, 2). Mammalian complex I, one of the largest membrane-bound enzymes in the cell, contains 45 subunits with a combined mass of 1 MDa; the fourteen fully-conserved core subunits are required for catalysis, while the 31 supernumerary subunits may be required for enzyme assembly, stability or regulation (3–8). The ‘active-deactive’ transition of mammalian complex I has recently come to prominence as a physiologically-relevant mechanism of regulation. In the absence of substrates, complex I relaxes into a profound resting state, known as the deactive state, that can be reactivated by addition of NADH and ubiquinone (9–12). Notably, because the respiratory chain cannot catalyze in the absence of O_2_, ischemia promotes complex I deactivation (13–15). Forming the deactive state may be protective because, upon reintroduction of O_2_ to the ischemic tissue, it is unable to catalyze the reverse electron transport reaction that causes a damaging ‘burst’ of reactive oxygen species production (16). Controlling complex I reactivation thus provides a rational strategy for combating ischemia-reperfusion injury (17, 18). Conversely, forming the deactive state may also tend to increase ischemia-reperfusion injury because it is more susceptible to oxidative damage than the active state (19), and strategies to target and protect the deactive state may also prove effective.

Rapid progress has been made recently in the structure of mammalian complex I due to a proliferation of structures for the *Bos taurus* (bovine) (5, 6, 20), *Sus scrofa* (porcine) (21, 22), and *Ovis aries* (ovine) (8, 23) enzymes, both in their isolated forms and in supercomplex assemblies. All 45 subunits of the mammalian complex have been assigned (5, 6, 20) and modeled (6, 8, 22), and in data from the bovine complex three different structural classes were identified (6). The three classes were tentatively assigned to different functional states of the complex. In the state referred to as class 1, several regions around the ubiquinone-binding site were disordered, whereas clear densities for them were observed in class 2. One of these regions is the loop between the first and second transmembrane helices (TMHs) of subunit ND3, which contains the reactive cysteine residue (Cys39) used as a biochemical marker for the deactive state (24). Cys39 can only be modified with thiol-reactive reagents such as *N*-ethylmaleimide (NEM) in the deactive state (24, 25). Because the cysteine is occluded in class 2, but likely more accessible on its unstructured loop in class 1, class 1 was tentatively assigned to the deactive state, and class 2 to the active state (6). A less populated class that is most similar to class 1, class 3, was also observed and refined to lower resolution. Its density map contains additional regions of disorder, including in part of the transverse helix that runs along the membrane domain appearing to strap it together, and so class 3 was ascribed to enzyme molecules in the process of dissociation (6).

These tentative assignments of the active and deactive structures suggest that the deactive state results when structural elements around the ubiquinone-binding site, including loops in the ND1, ND3 and 49 kDa subunits, become disordered (6). It has been proposed that the localized disorder disrupts the substrate binding site (rendering it catalytically inactive), but that ubiquinone interacting with the site serves as a template to re-structure it (the deactive state slowly reactivates when NADH and ubiquinone are provided) (6, 26). We refer to this model as the ‘unfolded Q-site’ model. Alternatively, other researchers have proposed the ‘truncated Q-site’ model. In the crystal structure of complex I from *Yarrowia lipolytica* (27), considered to be in the deactive state, the top of the ubiquinone-binding cavity is occluded by the β1-β2 loop of the 49 kDa subunit, preventing the ubiquinone headgroup reaching its binding site. The β1-β2 loop was subsequently modeled in a similar configuration in the structure of ovine complex I (8), and this structure also ascribed to the deactive enzyme.

Here, to define the structure of the deactive state we prepared biochemically-defined samples of deactive bovine complex I and determined their structure by single particle electron cryo-microscopy (cryoEM). Our preparations exhibit the well-known biochemical characteristics of the deactive state of the mammalian enzyme (9–11, 24), and are highly catalytically active following reactivation. The structure of the deactive complex matches the previously-described class 1 structure (6) and supports the unfolded Q-site model for the deactive transition. Thus, our model provides a structural foundation for interpreting the wealth of mechanistic, biochemical and physiological data on the deactive transition in mammalian complex I, and for understanding the role of deactive complex I in ischemia-reperfusion injury.

## Results

### Preparation of highly-active complex I in the deactive state

First, we developed a protocol to purify highly-active bovine complex I set fully in the deactive state. Our method was developed from that of Jones et al. (28), but with the final gel filtration step performed in the detergent cymal-7, rather than in n-dodecyl β-D-maltoside (DDM), as it was observed previously that cymal-7 gave a higher density of particles on cryo-EM grids (5). To convert the complex to the deactive state, the suspension of mitochondrial membranes from which the preparation begins was incubated at 37 °C for 15 minutes, before the detergent was added for solubilization. The temperature and length of incubation were optimized by testing the exposure of Cys39 in the ND3 subunit with NEM (24). NEM only reacts with Cys39 when the complex is deactive, and it prevents reactivation, so the activities observed in the presence and absence of NEM can be used to quantify the deactive and active states. In the presence of NEM the purified deactive enzyme displayed a very slow, constant rate of catalysis, whereas in its absence a pronounced lag phase was observed as the enzyme slowly reactivated. The maximal rate of catalysis was ~20 times higher in the absence of NEM than in its presence, indicating that the complex was ~95% in the deactive state (see Figure 1). Finally, by paying particular attention to the time taken for each stage of the preparation, the specific activity of the enzyme imaged here (following reactivation) was improved from the value described previously (14 ± 3 µmol NADH min^-1^ mg^-1^) (28), to 22.2 to 24.7 µmol NADH min^-1^ mg^-1^ (~390 NADH s^-1^). The activities of equivalent preparations carried out without the deactivation step were comparable. These activities match those of the mammalian complex in its native membrane (28) and are similar to the highest activities reported for isolated bacterial complex I (29, 30), so they confirm both the integrity of the purified complex and the reversibility of the deactivation procedure.

**Figure 1.**
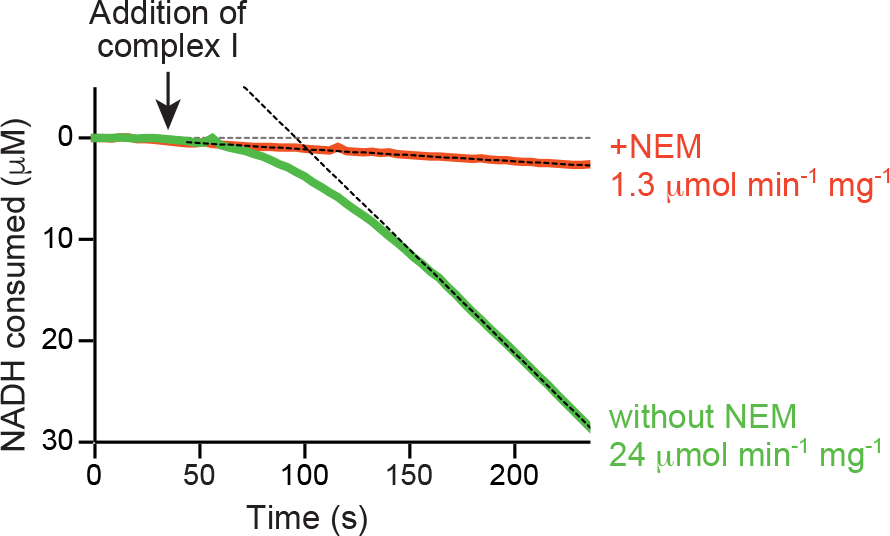
Spectrophotometric catalytic activity assay of NADH:decylubiquinone oxidoreduction by isolated deactive complex I. Assay traces comparing enzyme that had been treated by 4 mM NEM (red) with enzyme that had not been treated (green). Without the NEM treatment the deactive protein gradually reactivates, reaching its maximal rate after 150 s. The NEM treatment prevents reactivation and the background rate is only from the small proportion of active enzyme present. Experiments were carried out using 200 µM NADH, 200 µM decylubiquinone and 0.5µg mL^-1^ complex I, as described in Methods.

### Imaging, classification and structure modeling for the deactive enzyme

Quantifoil holey carbon grids were used previously to image mammalian complex I (5, 6, 8). However, in common with many other proteins, the complex binds to the oxidatively-modified carbon, depleting it from the vitreous ice in the holes and leading to poorly distributed particles and low particles number for imaging. Self-assembled monolayers with controlled surface properties and lower protein affinities have been developed to mitigate this problem (31), and here we used UltrAuFoil gold grids (32) derivatized with a polyethylene glycol (PEG) linked by an alkanethiol; the 11-carbon alkanethiol forms a robust bond to the gold surface, and exposes the biocompatible PEG-6 group to the protein solution (31). In a side-by-side comparison with the previously-used Quantifoil grids, we found four times more particles could be imaged per hole using the PEGylated gold grids (see Figure 2), plus the particle distribution was improved and less aggregation observed (see Figure S1). Subsequently, it also became clear that the particles adopt a broader set of orientations on the PEGylated gold than the Quantifoil grids (see Figure S2). In addition to altered grid-protein interactions, varying ice thicknesses may also contribute to the improved distribution, and the amorphous carbon Quantifoil grids may absorb detergent from the solution, altering the properties of the air-water interface during grid preparation and increasing the chance of aggregation.

**Figure 2.**
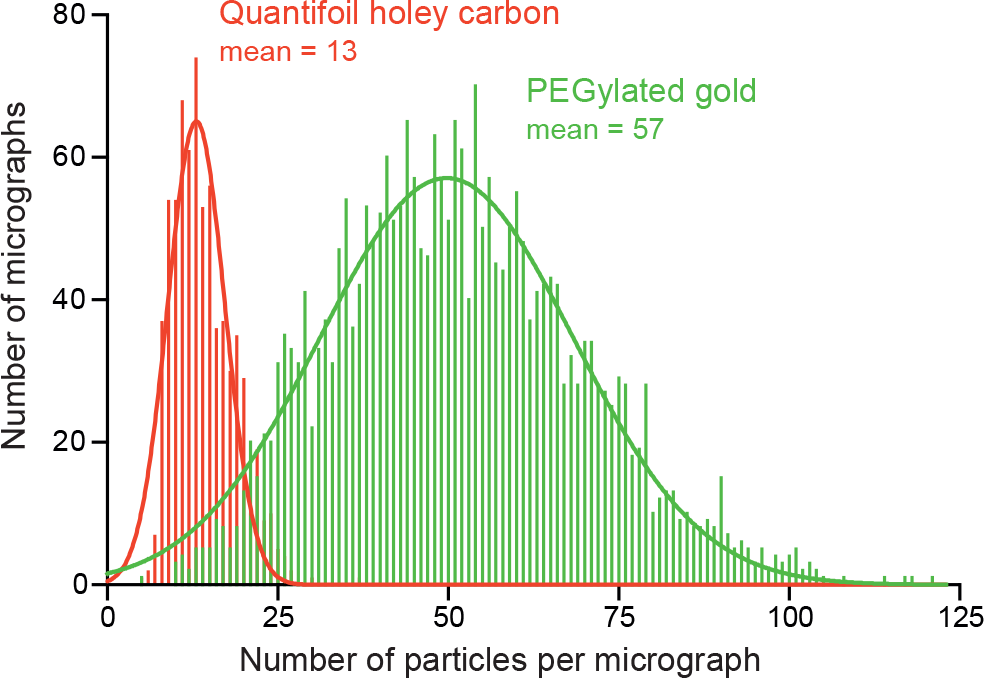
Comparison of the number of particles observed per micrograph using PEGylated gold and Quantifoil holey carbon grids. The samples of deactive complex I used were at concentrations of 4.4 mg mL^-1^ (PEGylated gold UltrAuFoil 0.6/1) and 4.2 mg mL^-1^ (Quantifoil 0.6/1). The PEGylated gold grids were prepared using a Vitrobot (see Methods) and the Quantifoil grids by manual blotting as described previously (5, 6). The data are from two automated data collection sessions on a Titan Krios microscope (see Methods for imaging parameters) and the particles were picked manually in each case.

The deactive complex on the PEGylated gold grids was imaged at 300 kV using a Titan Krios electron microscope and a Falcon-II direct electron detector (33). A total of 148k particles were picked manually, and 125k particles were retained following 2D and coarsely-sampled 3D classification. Using the RELION software suite (34, 35), the data set was first refined to produce a 4.7 Å resolution density map. Following per-particle frame alignment and B-factor weighting (35), the final resolution was 4.13 Å (defined where the Fourier shell correlation (FSC) = 0.143) (36) (see Figure S3).

The 125k particles with improved signal to noise following frame alignment and B-factor weighting were then subjected to 3D classification with incrementally increasing angular sampling (up to 0.9°) (37). The results are shown in Figure 3. Classification into three classes resulted in a dominant class containing 87.5% of the particles, a minor class containing 9.7%, and a negligible third class containing 2.7%. When the classification was repeated but with six classes a similar pattern emerged: two major classes contained 87.0% and 7.9% of the particles (matching their equivalent classes from before) and the remaining four classes were all negligible (1.3%, 1.6%, 0.6%, and 1.5%). The two largest classes from the first evaluation were refined individually, leading to cryoEM density maps of formally 4.13 and 7.50 Å resolution. The map for the dominant class at 4.13 Å resolution, which we assign to the structure of deactive complex I, was taken forward to model building (see Table S1). Although the formal resolution of 4.13 Å is only marginally higher than reported previously for the bovine complex (4.27 Å for class 1 and 4.35 Å for class 2) (6), several regions of the map displayed substantially improved features.

**Figure 3.**
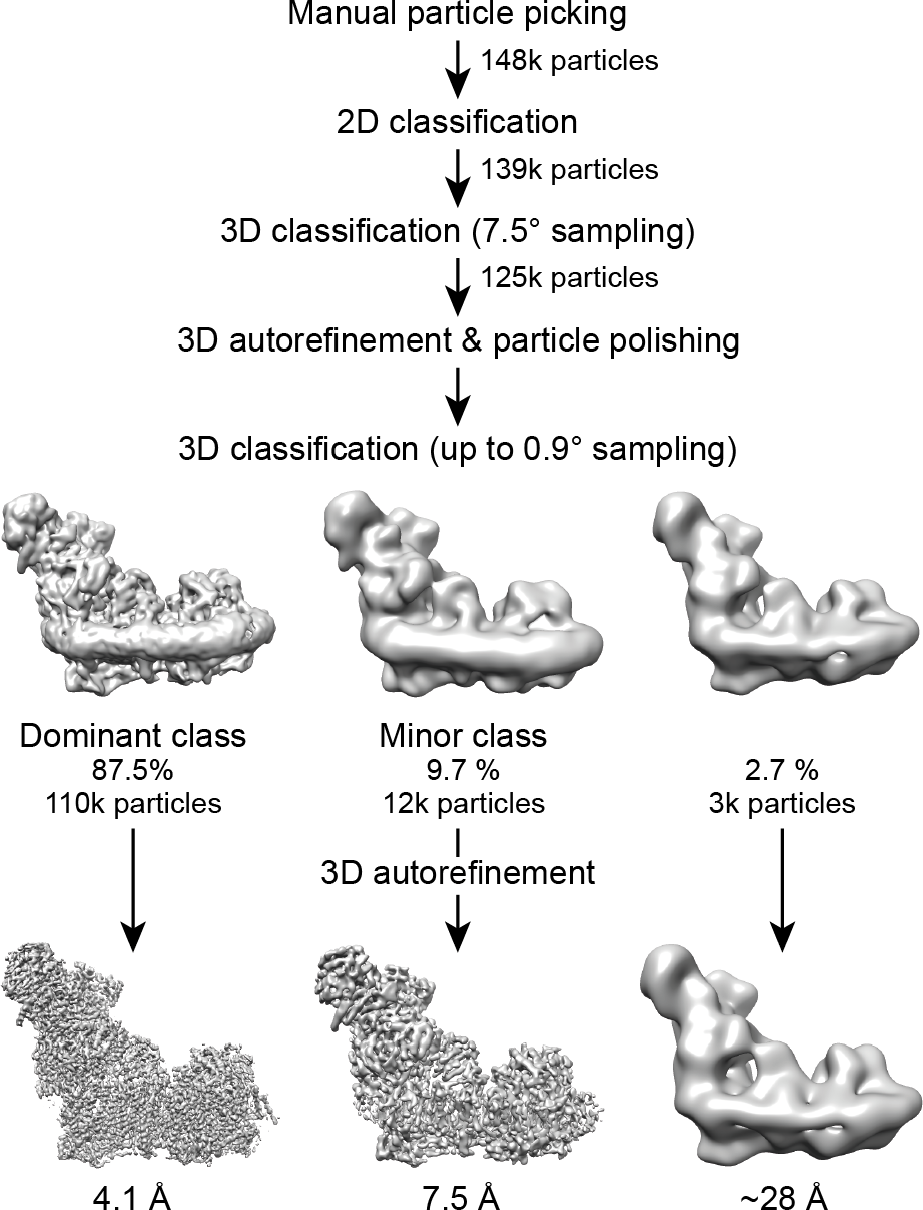
Classification and refinement of the cryoEM density map for deactive complex I. The RELION pipeline (34) was used to process data from the deactive preparation. Following manual particle picking and 2D and 3D classification to discard bad particles, 3D refinement and particle polishing were performed. Subsequently, the particles were classified using an angular sampling up to 0.9° and with the resolution limited to 8 Å (see Methods). All the three classes provided were populated. The dominant class refined to 4.1 Å, and a minor class to 7.5 Å. The remaining class is negligible as it contained so few particles.

Consequently, by incorporating information from the recently published ovine model and map (8) we were able to assign sequence to the large domain of the 75 kDa (NDUFS1) core subunit (see Table S2), to provide a fully-assigned model for all 14 core subunits (note that we refer to the subunits using both their bovine and human nomenclatures as summarized in Tables S2 and S3). In addition, we were able to fully assign the sequences of the 42 kDa (NDUFA10), 18 kDa (NDUFS4), 13 kDa (NDUFS6), 10 kDa (NDUFV3), PGIV (NDUFA8), SGDH (NDUFB5), and B22 (NDUFB9) supernumerary subunits, and to increase the level of sequence assignment in the B16.6 (NDUFA13), B15 (NDUFB4), B14.5a (NDUFA7), and B14.5b (NDUFC2) subunits (see Table S3). Overall, our model contains 7,811 residues, of which 7,004 (90%) are assigned, increased from 71% in the previous class 1 model for the bovine enzyme (6).

### Structural rearrangements of the ubiquinone-binding site region in deactive complex I

The cryoEM density map for the biochemically-defined deactive enzyme contains specific regions of localized disorder around the ubiquinone-binding site. Continuous densities for the loop between TMHs 5 and 6 in the ND1 subunit, the loop between TMHs 1 and 2 in the ND3 subunit, the short loop between the β1 and β2 strands in the 49 kDa (NDUFS2) subunit, and several nearby loops in the 39 kDa (NDUFA9) subunit are not observed in the map (see Figure 4 and Figure S4). The same regions are absent from the previously-described class 1 density map, but were clearly observed in the class 2 map, despite its resolution being lower (6). Therefore, the loss of ordered structures around the ubiquinone-binding site is characteristic of the deactive enzyme. The ubiquinone access channel was tentatively identified in the class 2 map, leading from an entrance in ND1 to the binding site for the ubiquinone headgroup, at the top of a cleft between the 49 kDa and PSST subunits (6) (see Figure 4). A similar channel was identified in the crystal structure of complex I from *Thermus thermophilus*, which does not exhibit a clear active-deactive transition and can thus be assumed to be in an active state (38). The channel cannot be identified in the structure of the deactive enzyme because key structural elements that form it are disordered and missing from the model. We conclude that the ubiquinone-binding channel has lost its structural integrity in the deactive complex.

**Figure 4.**
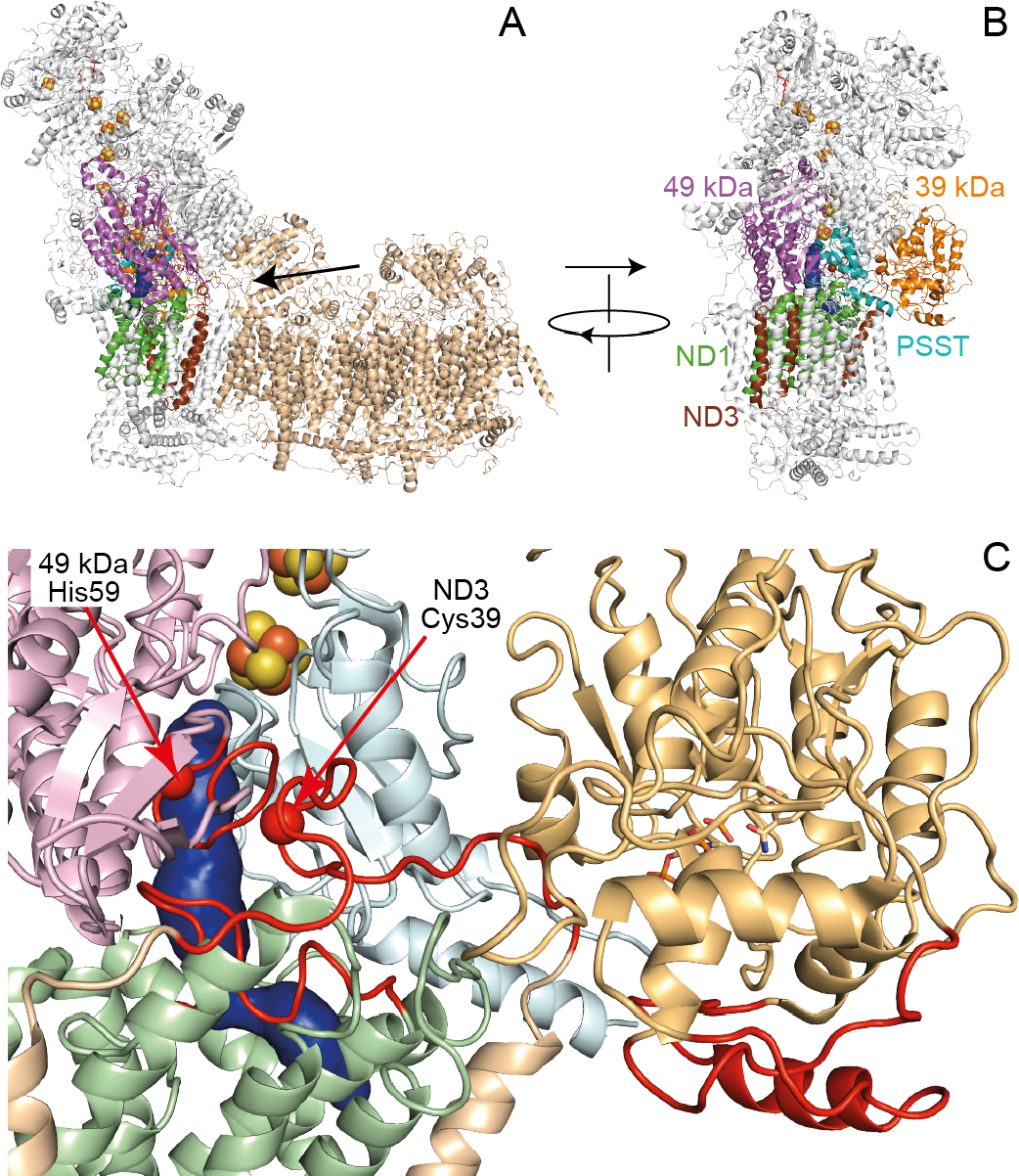
The structure of deactive complex I is characterized by localized unfolding. A) Structure of intact complex I with an arrow showing the view taken of the ubiquinone-binding region. The distal section of the membrane domain (shown in wheat) is not included in panels B and C. B) View of the ubiquinone-binding region with the subunits involved shown in color as indicated. C) Close up of the ubiquinone-binding region (same perspective as in B) with the ubiquinone-binding channel predicted for the class 2 structure (6) shown in blue. The colors are lighter colored versions of those used in B. The areas that become disordered in the deactive state (the loops between TMHs 1 and 2 in ND3, TMHs 5 and 6 in ND1, ²1 and ² 2 in the 49 kDa (NDUFS2) subunit, and parts of the 39 kDa (NDUFS9) subunit) are shown in red. His59 is one of the residues likely to interact with the bound ubiquinone head group, Cys39 is the marker residue for the deactive state. The figure was created by combining 5LC5.pdb for the active enzyme (6) with information about the deactive state described here.

### Assignment of classes 1, 2 and 3

The preparation of bovine complex I imaged by Zhu and coworkers (6) comprised a mixture of deactive and active enzymes and three structural classes (classes 1, 2 and 3) were distinguished. Disordered regions observed here in the biochemically-defined deactive complex were disordered in class 1, but not in class 2. The 5.6 Å resolution of the class 3 density map is too low for a similar comparison, but class 3 is clearly distinguished by additional disorder (not present in classes 1 and 2) in the C-terminal section of ND5 (including the transverse helix and TMH16) and part of the adjacent subunit B14.7 (NDUFA11), consistent with it being partially dissociated and irreversibly inactivated (6). ND5 and B14.7 (NDUFA11) are both represented by clear density in the deactive enzyme (see Figure S4), supporting assignment of class 1 to the deactive state. Two global comparisons were further used to compare the deactive enzyme to classes 1, 2 and 3. Previously, small shifts and rotations in different enzyme domains were observed between the classes. For example, with the structures superimposed on ND1 (in the ‘heel’ of the enzyme), the hydrophilic domain rotates by 3.4° and the membrane domain by 3.9° between classes 1 and 2, and the distal portion of the membrane domain a further 3.1° between classes 1 and 3 (6). To capture these global rearrangements, map/model correlations were calculated to evaluate how well the Cα chains from classes 1, 2 and 3 fit the deactive complex I density map (see Table 1).

In addition, root-mean-squared-deviation (RMSD) values were obtained for different sections of the membrane domain, following alignment of the hydrophilic domains of the deactive model and classes 1, 2 and 3 (see Table 1). Both approaches support the assignment of class 1 to the deactive enzyme. Furthermore, the minor (9.7%) class in the deactive preparation matches the class 3 structure reported previously (6); it displays disorder in both the C-terminal section of ND5 and subunit B14.7 (NDUFA11) and has the highest map/model correlation with class 3 (see Table 1). This match is consistent with a small proportion of partially-denatured enzyme in grids of the deactive complex. With class 1 confirmed as the deactive state, increased structuring of the ubiquinone-binding site in class 2, and the similarities between the ubiquinone-binding channels detected in class 2 and in complex I from *T. thermophiles* (6, 38) argue strongly that class 2 represents the active enzyme, the state that is ready for substrate binding.

**Table 1-.** Comparison of the structure of the deactive enzyme with the previously-determined class 1, 2 and 3 structures. Map/model correlations were from UCSF Chimera and obtained by fitting the Cα chains for classes 1, 2 and 3 into the maps from the deactive preparation, where larger values indicate a better fit. RMSD values are for the Cα coordinates for the deactive complex (dominant class) compared to classes 1, 2 and 3, where smaller values indicate a better fit. Following superposition (using the Pymol ‘super’ routine) of all the models on the Cα structures of the core subunits in their hydrophilic domains (chains B, C, D, E, F, G and I), RMSD values were calculated for sets of core subunits in the membrane domain. The core hydrophobic domain consists of chains A, H, J, K, L, M and N, the proximal domain of chains A, H, J, K and N, and the distal domain of chains L and M. The N-terminus (residues 1-39) of chain D was excluded. A model for the minor class was generated by performing real-space rigid-body fitting of each of the dominant class model chains to the minor class density in Phenix (59). Values that match best to the different classes are in bold and marked with asterisks.

|  | **Map/model correlation** |  |  |
| --- | --- | --- | --- |
|  | Class 1<br>5ldw.pdb | Class 2<br>5lc5.pdb | Class 3<br>5ldx.pdb |
| Dominant class (this study) | **0.1988\*** | 0.1496 | 0.1772 |
| Minor class (this study) | 0.1097 | 0.1026 | **0.1218\*** |
| Ovine-brij (EMD-4084) | 0.1615 | 0.1388 | **0.1775\*** |
| Ovine supercomplex (EMD-8130) | **0.1376\*** | 0.1206 | 0.1285 |
| Porcine supercomplex (EMD-9539) | 0.0718 | **0.0745\*** | 0.0635 |
|  | **RMSD values (dominant class)** |  |  |
| Hydrophilic domain | 1.75 | 1.77 | 1.77 |
| Hydrophobic domain - all | **1.33\*** | 5.88 | 3.16 |
| Hydrophobic domain - proximal | **0.90\*** | 3.55 | 1.03 |
| Hydrophobic domain - distal | **1.63\*** | 7.42 | 4.28 |
|  | **RMSD values (minor class)** |  |  |
| Hydrophilic domain | 1.78 | 1.80 | 1.77 |
| Hydrophobic domain - all | 5.33 | 7.37 | **1.94\*** |
| Hydrophobic domain - proximal | 1.70 | 3.89 | **0.94\*** |
| Hydrophobic domain - distal | 7.23 | 9.53 | **2.54\*** |

### Comparison with structures from other mammalian species

Using similar criteria, the classes represented by published cryoEM structures from other mammalian species were evaluated. For the isolated ovine complex in brij-35 at 3.9 Å resolution (8), no continuous densities are present for the loops that are disordered in the bovine deactive state, or for the C-terminal section of ND5 or subunit B14.7 (NDUFA11) that are disordered in bovine class 3 (see Figure S4). RMSD calculations of the protein models between different species are confounded by variations in sequence numbering, but map/structure correlations (see Table 1) support the ovine structure matching the bovine class 3 (inactive) state. Previously, the ovine structure was assigned to the deactive state (8), and a second, lower-resolution class (not available for analysis) to the active state. However, the specific activity of the preparation (in the detergent brij-35 to increase particle density on the grids) was only 3.5 µmol NADH min^-1^ mg^-1^ (8, 39), equivalent to ~16% of the activity reported here and consistent with a large proportion of inactive enzyme. For complex I in the ovine supercomplex in digitonin (23) the resolution is too low to observe specific loops, but map/structure correlations suggest it is predominantly in the class 1 (deactive) state. For complex I in the porcine supercomplex, also in digitonin (21, 22), map/structure correlations indicate it is in the class 2 (active) state and this is supported by clear density for the ND3, ND1, 49 kDa (NDUFS2) and 39 kDa (NDUFA9) subunit loops (see Figure S4). A recent structure of the bovine supercomplex is too low resolution to allow either specific structural elements or map/model correlations to distinguish the class (40). Finally, although supercomplex formation is independent of the deactive/active status of complex I (41), incorporating complex I into a supercomplex may both stabilize the unstable region around the C-terminus of ND5 and subunit B14.7 (NDUFA11) that is buttressed against complex III, and influence the conformation of the membrane arm as it curves around it.

## Discussion

The complex I preparations used here have specific activities at least two-fold higher than those used in our previous cryoEM studies of bovine complex I (6, 42). However, based on particle classification (37), the fraction of class 3 inactive particles has only decreased from ~20% to ~10%, questioning whether the class populations observed on the grids accurately reflect the class populations in solution. Many proteins denature at the large air-water interface present during cryoEM grid formation (43), so class 1 and 2 molecules may augment the class 3 population, or become more completely denatured and (having lost their distinctive L-shape) invisible to the analysis. Our observation cautions against relying on the classification of mixed populations of subtly-different particles when assigning biochemically known states, and suggests that higher resolution structures of mammalian complex I set in catalytically-relevant states will require homogeneous preparations combined with solution conditions that maintain their stability during grid preparation. Here, the deactive complex was prepared in a homogeneous state that is reflected in the class populations on the cryoEM grids, giving confidence in its structural assignment.

The structure of bovine complex I set in the deactive state supports the unfolded Q-site model (6, 26) for the deactive transition of the mammalian enzyme. The competing truncated Q-site model was originally proposed using structural data from *Y. lipolytica* complex I (27). However, the structure described has an inhibitor bound adjacent to the loop that truncates the channel, which may stabilize it in an alternative conformation. Furthermore, the transition is less pronounced in *Y. lipolytica* than in mammalian species: interconversions between the active and deactive states are faster and associated with much lower activation energies (44, 45). Thus, the structural changes of the deactive transition in the yeast enzyme may be less extensive than in the mammalian enzyme. The ubiquinone-binding channel was observed to be similarly truncated in the structure of ovine complex (8). However, the ovine complex has very low specific activity, correlates structurally to the inactive class 3 bovine complex, and the density for the loop in question is not well resolved (see Figure S4). The ovine structure (and bovine class 3) probably represent inactive states that cannot be reactivated.

The flexibility of the structural elements that become disordered in the deactive state (see Figure 4) is further underlined by the different conformations they adopt in structural models determined for different species, from the mammalian (8, 22), yeast (27), and bacterial (38) enzymes, consistent with them having important functional roles. The ND3 loop appears to be a ‘tether’ from the membrane domain, on the front of the hydrophilic domain and ubiquinone-binding channel; the ND1 loop forms the base of the ubiquinone-binding channel at the hydrophobic-hydrophilic domain interface; the β1-β2 loop in the 49 kDa (NDUFS2) subunit carries a histidine that ligates the ubiquinone headgroup. All these loops are crucial for both the integrity of the ubiquinone-binding channel, and the structure of the domain interface, which they appear to maintain in an activated state (analagous to a compressed spring) in the active enzyme. Upon deactivation the interface relaxes, with consequent changes to the relative arrangement of the two domains. Thus, we propose that the deactive state is a reversibly-formed off-pathway state and not, as suggested previously (27), a catalytic intermediate. The disordered elements are confined by adjacent secondary structures, and the disordered region in general may be stabilized by the supernumerary 39 kDa (NDUFA9) subunit, on the outside of the core complex. In the inactive class 3 structure, loss of structural integrity in the ND5 transverse helix appears to allow further relaxation within the membrane domain, and the proximal section of the membrane domain to begin to break from the rest of the complex (6). Thus, like the loop in ND3, the transverse helix can also be considered to be a tether that maintains the enzyme in an active conformation.

Structural knowledge of the deactive state of mammalian complex I now provides a basis for understanding many of its biochemical features. i) Cys39 in subunit ND3, which is both used as a marker for the deactive state (24) and targeted in strategies to minimize ischemia-reperfusion injury, by using cysteine-modifying agents to slow reactivation or to protect the cysteine against irreversible oxidation (15, 17, 19), is on the (disordered) loop between TMHs 1 and 2. It is occluded in the active state and must become solvent accessible in the deactive state. ii) Structural disorder in the ND3, ND1 and 39 kDa (NDUFA9) subunits in the deactive state explain the results of cross-linking studies that identified these subunits as changing conformation in the deactive state (41, 46). iii) Relaxation of the activated interface between the hydrophobic and hydrophilic domains upon formation of the deactive state is consistent with the functional connection between them breaking down upon deactivation. Thus, the proton transfer subunits in the hydrophobic domain are freed from control by the redox reaction in the hydrophilic domain and may function independently of it, resulting in the Na^+^/H^+^ antiporter activity that has been observed specifically in the deactive state (47). iv) The unfolded Q-site model for the deactive state explains why slow reactivation of the deactive enzyme only occurs in the presence of NADH and ubiquinone (9). We propose that ubiquinone acts as a template to restructure the site in the NADH-reduced enzyme, in an induced-fit mechanism of substrate binding (48). The requirement for ubiquinone explains why neither reverse electron transfer (ubiquinol:NAD^+^ oxidoreduction) nor its associated reactive species production are catalyzed by the deactive enzyme upon the reperfusion of ischemic tissue (9, 17). v) The ubiquinone-site inhibitor rotenone has also been reported to return the deactive enzyme to its active state (49). consistent with its inhibition of the Na^+^/H^+^ antiporter activity of the deactive state (47). vi) The flexibility and ability of the structural elements that constitute the active site to reorganize around substrates and inhibitors may explain why so many diverse compounds are known to inhibit ubiquinone reduction by complex I (50, 51). Similarly, the instability of the ubiquinone-binding site region, which propagates structural flexibility through the enzyme, may explain why the mammalian enzyme has proved so difficult to purify in a highly-catalytically active state and (so far) to crystallize for structure determination.

Finally, disordered protein domains are increasingly recognized as central to many diverse molecular processes and as particularly important in regulatory mechanisms (52). The deactive state of complex I is already being explored as a regulatory mechanism relevant to minimizing ischemia-reperfusion injury (17), and the structure of the deactive state now highlights additional possibilities. Inherent conformational flexibility in the loop of ND3 that carries the highly-conserved Cys39 may transiently expose it to post-translational modifications that regulate complex I activity in response to cellular redox status. Alternatively, several disordered regions accumulated into one area may allow an effector protein to interact, to trap the enzyme in the deactive state, or promote its formation. Studies of the deactive-active status of complex I under physiologically-relevant conditions and of the deactive state formed *in vivo* will be required to investigate these suggestions in the future.

## Materials and methods

### Preparation of complex I samples

Bovine mitochondria and mitochondrial membranes were prepared as described previously (53). Complex I was placed in the deactive state by incubating the membranes (resuspended to 12 mg protein mL^-1^ in 20 mM Tris-Cl pH 7.55, 1 mM EDTA, 10% glycerol, 0.0075% PMSF) at 37 °C for 15 min. Then, the membranes were diluted to 5 mg mL^-1^ in the same buffer but ice cold, and cooled on ice for 10 min. All subsequent steps were performed at 4 °C, using a protocol developed from that of Jones et al. (28). Briefly, *n*-dodecyl β-D-maltoside (DDM, Glycon Biochemicals GmbH) was added dropwise to 1%, the suspension stirred for 20 min., clarified by centrifugation (47,000 × g for 12 min.) and loaded onto a Q-sepharose column pre-equilibrated in buffer A (20 mM Tris-Cl pH 7.55, 2 mM EDTA, 10% ethylene glycol, 0.2% DDM, 0.02% asolectin (Avanti Polar Lipids) and 0.02% CHAPS (Santa Cruz Biotechnology)). Cytochrome *c* oxidase and other unwanted proteins were eluted in 27.5% buffer B (buffer A with 1 M NaCl added), until the absorbance at 420 nm reached 0.025, then complex I was eluted in 36% buffer B. The complex I-containing fractions were pooled and concentrated to ~1 mL, then eluted from a 10/300 superose-6 increase column (GE Healthcare Life Sciences) at 0.5 mL min^-1^ in 20 mM Tris-Cl pH 7.55, 150 mM NaCl, and 0.04% Cymal-7 (Anatrace). The manually collected peak fraction with concentration ~4 mg mL^-1^ was used immediately for grid preparation.

### Catalytic activity assays and determination of the deactive/active enzyme ratio

NADH:decylubiquinone oxidoreductase activities of isolated complex I samples were determined at 32 °C using 0.5 µg complex I mL^-1^ with 200 µM NADH and 200 µM decylubiquinone in 20 mM Tris-Cl pH 7.5, 0.15% asolectin and 0.15% CHAPS. The reaction was initiated by addition of NADH and the rate determined following activation of the deactive enzyme, when (typically 2 min. after initiation) the kinetic trace (recorded at 340 −380 nm, ε_NADH_ = 4.81 mM^-1^ cm^−1^) becomes linear. To determine the deactive/active enzyme status an aliquot of the complex I stock solution (at ~4 mg mL^−1^) was divided into two and 4 mM NEM added to one half. The samples were incubated at 4 °C for at least 5 minutes (longer incubations did not increase the level of inhibition) before their addition to the assay mixture.

### Cryo-EM grid preparation

For the data collection presented, UltrAuFoil gold grids (0.6/1, Quantifoil Micro Tools GmbH) (32) were glow discharged at 20 mA for 60 s then imported to an anaerobic glovebox and placed in ethanol containing 5 mM 11-mercaptoundecyl hexaethyleneglycol (SPT-0011P6, SensoPath Technologies) for at least 24 hours before grid preparation (31). Then, just prior to use, the grids were washed three times in ethanol and left to air-dry. Grids were prepared using an FEI Vitrobot IV. 2.5 µL of protein solution were applied to the grid at 4 °C in 100% relative humidity, and blotted for 8 – 12 s at force setting −10, before being plunged into liquid ethane. For comparative experiments, UltrAuFoil 1.2/1.3 gold grids and Quantifoil 0.6/1 grids were prepared similarly, but using 8 and 2 s blotting times, respectively, or prepared by manual blotting as described previously (5).

### Electron microscopy

Grids were imaged in a 300 keV Titan Krios microscope fitted with a Falcon-II direct electron detector and EPU software at the Electron Bio-Imaging Centre at The Diamond Light Source. The nominal magnification was set to 59,000× but, by comparing the final 4.13 Å structure to the previously published class 1 structure, the pixel size was calibrated to 1.38 Å and the magnification to 101,449×. A C2 and objective aperture of 100 µm were used and each image was exposed for 2.5 s with a total dose of ~80 electrons/Å^2^. We collected the first 12 frames (700 ms) to capture the rapid early movement of the sample when the electron beams first interacts with the grid (54) and after that every four frames were binned together. The defocus range was 1.3-3.1 µm in 0.3 µm increments; defocus was measured in the autofocus routine every 10 µm.

### Image processing

Whole-frame alignment was performing using Unblur (55) before CTF estimation using CTFFIND4 (56). All resolution estimates are based on the FSC = 0.143 criterion, and the final resolution estimates were made after the application of a binary mask and phase-randomization to check the effects of the mask. RELION-1.4 was used for data processing (34).

A total of 148,488 particles were picked manually. Following 2D and 3D classification to remove ‘bad’ particles, 125,006 particles were used for refinement to a resolution of 4.7 Å. Per-particle frame alignment to correct for movement and B-factor weighting (35) were then performed and the resolution, following a second refinement, improved to 4.13 Å with an angular accuracy of 0.87°. The resulting ‘shiny’ particles were subjected to 3D classification with three classes, with the angular sampling gradually increased up to 0.9° and the resolution limited to 8 Å to reduce over-fitting; local searches were implemented from 3.75° onwards. The populations within the classes remained stable for at least 50 further iterations after the different classes had emerged. The three classes (see Figure 3) were refined individually. The dominant class had a resolution of 4.13 Å and was sharpened with a B-factor of −110 Å^2^ before model building.

### Model building

We extended our previous class 1 model (6) to assign more residues, using the improved densities visible in the new map, and the better densities for the hydrophilic domain in the ovine map (8). Typically, the approximate numbering of the unknown residues (6) was found to be quite accurate, with bulky residues being found already placed in bulky pockets of density. In regions in which the numbering of the residues is still uncertain a poly-Ala chain was used to provide approximate numbers. The resulting model was further manually fitted in Coot (57) and refined using REFMAC5 (58). Sidechains were included where appropriate. Note that we number the residues in the subunits starting from the first residue of the mature protein (4). Note also that the density assigned to the 10 kDa (NDUFV3) subunit in both the bovine and ovine structures (6, 8) was assigned to the *N*-terminal mitochondrial targeting sequence of the 24 kDa subunit (NDUFV2) in the porcine structure (22) but it is cleaved from the mature protein and not present in the isolated enzyme (4).

## Acknowledgements

We thank A. Raine, M. Hartley and D. Gallagher (MBU) for computational help. Data were recorded at the UK National Electron Bio-Imaging Centre (eBIC) at Diamond (proposal EM13581, funded by the Wellcome Trust, MRC and BBSRC) with help from Dan Clare and Alistair Siebert. This work was supported by The Medical Research Council, grant numbers U105663141 to J.H. and U105184322 (K.R.V., in R. Henderson’s group).

---

**Figure S1.**
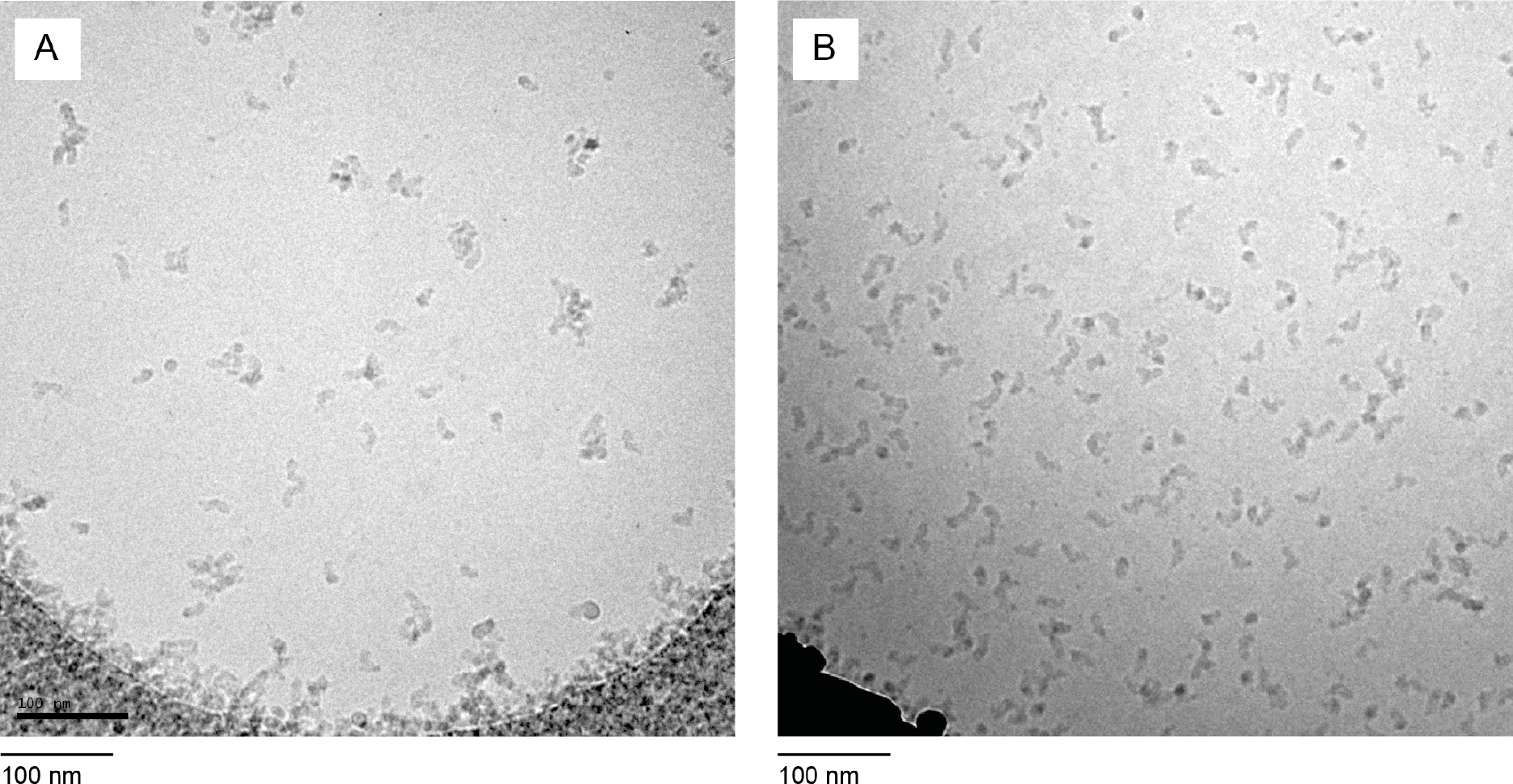
Micrographs from a sample of deactive complex I imaged on Quantifoil and PEGylated gold UltrAuFoil grids. A) Micrograph from a representative hole in a Quantifoil (0.6/1) grid that had been glow-discharged for 90 s at 20 mA. B) Micrograph from a representative hole in a PEGylated gold (1.2/1.3) grid prepared as described in Methods. Both grids were prepared using the same deactive complex I preparation, which had been frozen before grid preparation, using a Vitrobot (blot force −10, temperature 4 °C, and relative humidity 100%) but with longer blotting time (8 s) for the PEGylated gold grids than the Quantifoil grids (2 s) to account for their increased hydrophilicity. Both grids were imaged in an FEI T12 microscope with substantial defocus.

**Figure S2.**
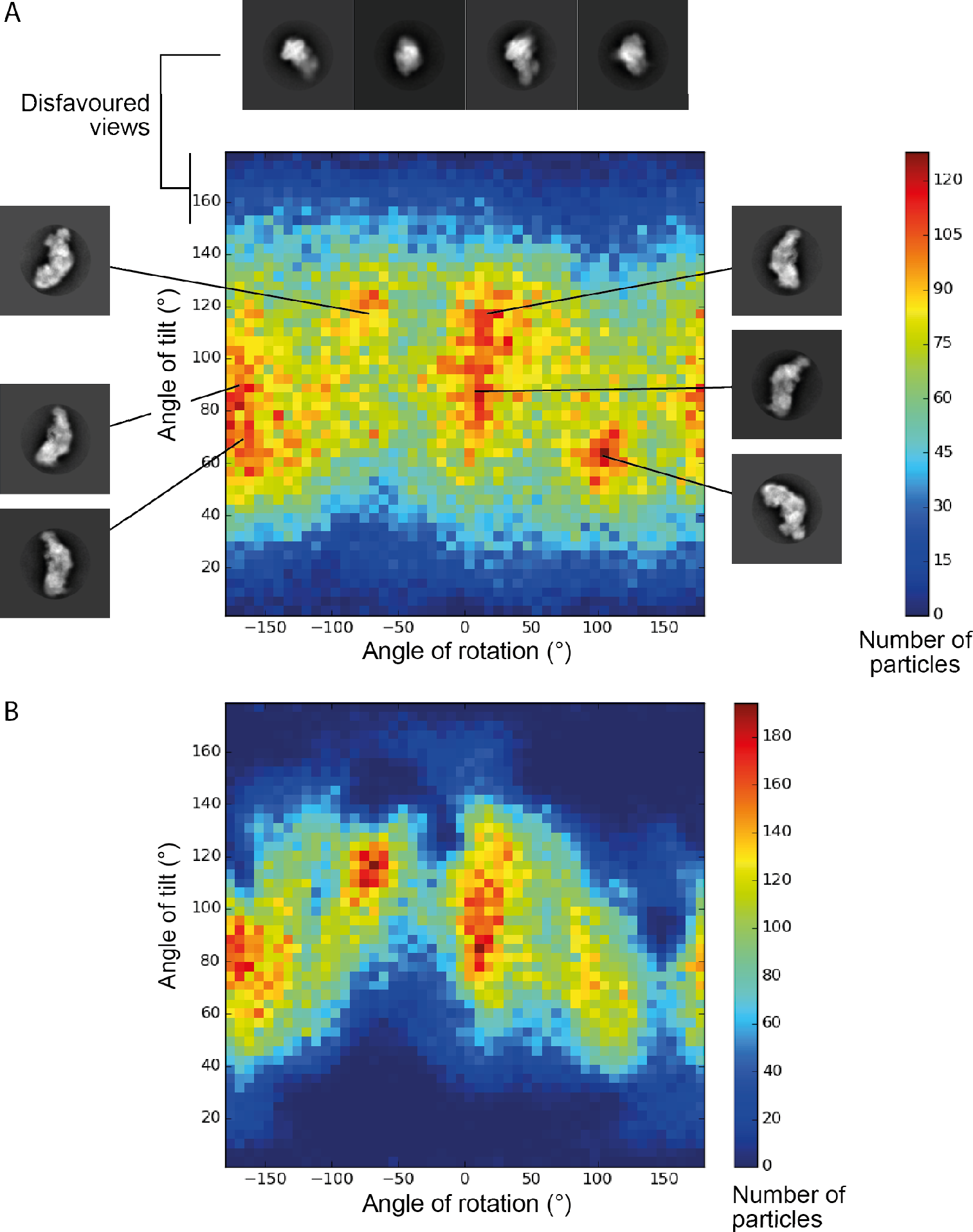
Analysis of the orientation of deactive complex I particles on PEGylated gold UltrAuFoil and Quantifoil grids. A) Distribution of particle orientations on PEGylated 0.6/1 gold grids with example 2D class averages from different regions of the plot. B) Particle orientations on 0.6/1 Quantifoil grids from a previously reported dataset (6). The PEGylated gold dataset contains 125,007 particles, and the Quantifoil dataset 115,974 particles; each dimension was split into 50 bins. The angles of rotation and tilt are taken from the ‘_rlnAngleRot’ and ‘_rlnAngleTilt’ values for each particle after autorefinement in RELION 1.4. As expected there was no correlation with the in-plane rotation (_rlnAnglePsi) parameter.

**Figure S3.**
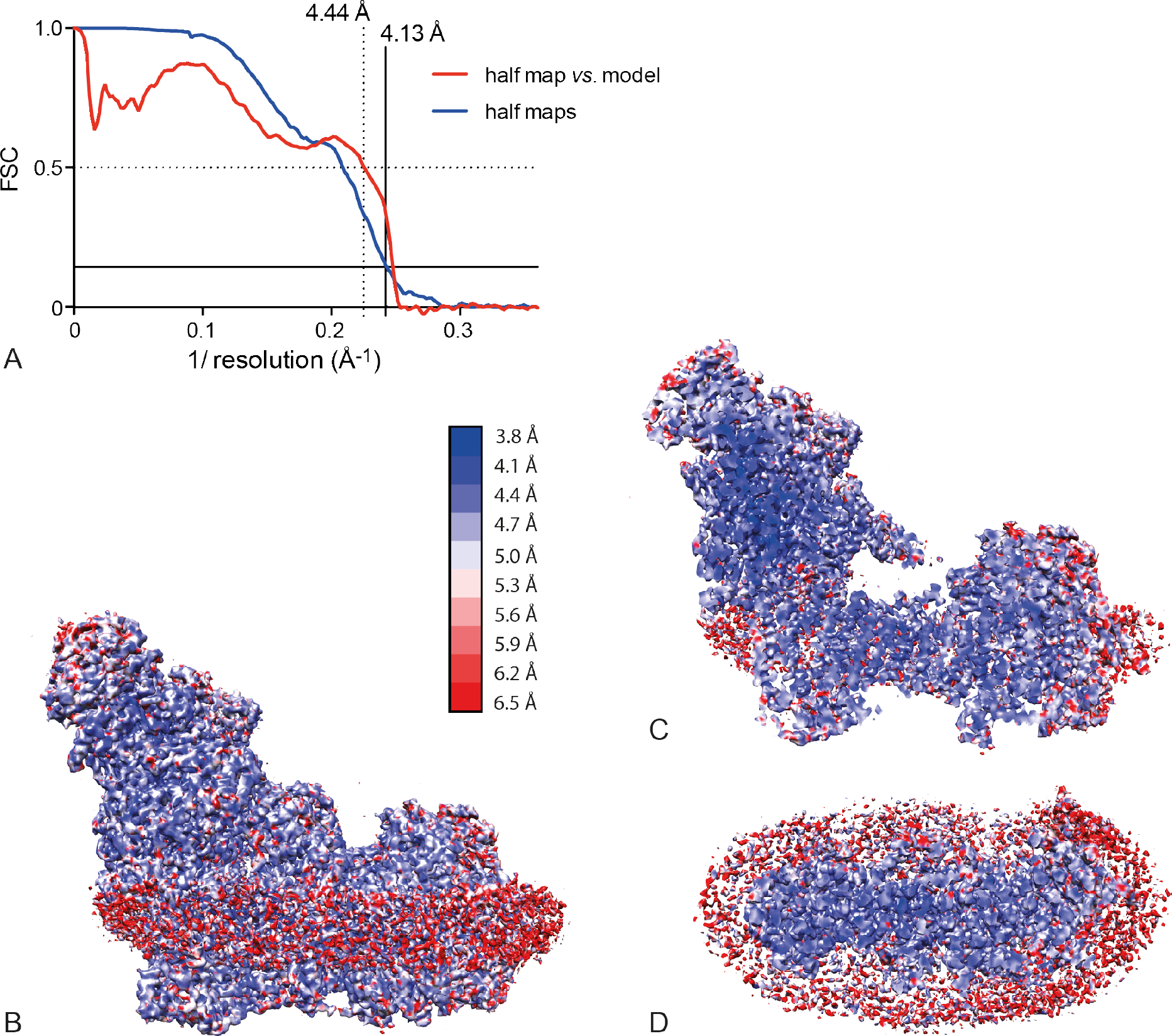
Resolution estimation and ResMap analysis for deactive complex I. A) FSC curve for the dominant class of the deactive preparation. B-D) Local resolution for the dominant class of the deactive preparation analyzed using ResMap (60). In panel C the resolution of the inner core of the complex, particularly around the iron-sulfur clusters, is shown to have a higher resolution than the outside (panel B); the amorphous detergent/phospholipid belt is poorly resolved. Panel D shows a cross-section of the complex from the matrix side, with the hydrophilic domain cut away.

**Figure S4.**
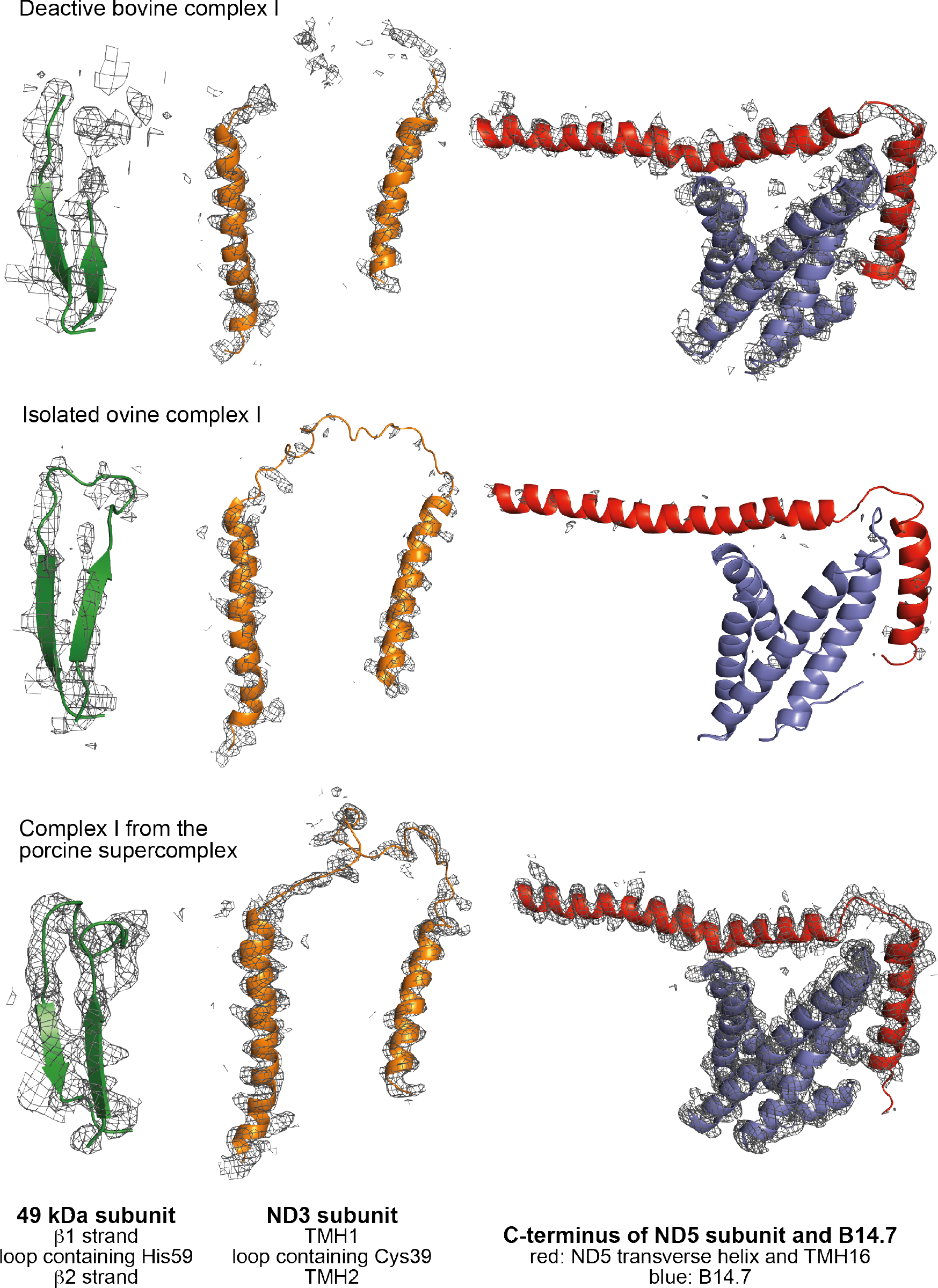
Comparison of characteristic densities in deactive bovine complex and in structures from other mammalian species. Green: comparison of densities for the β1-β2 loop in the 49 kDa (NDUFS2) subunit. Orange: comparison of densities for the TMH1-2 loop in subunit ND3. Red: comparison of densities for the C-terminus of subunit ND5 and blue: subunit B14.7 (NDUFA11). The contour levels for the density were set in Pymol to appropriate levels for the TMHs of subunit ND3, and then kept constant for the other subunits (deactive bovine complex I, 6.3; ovine complex I, 6.0; porcine supercomplex, 8.5). A carve radius of 2.5 Å was used throughout. The deactive bovine complex is from the data described here; the densities for elements not modeled were carved using the structures of the subunits from the active complex. The ovine complex I structure is for the isolated complex I of Sazanov and coworkers (8) (5LNK.pdb and EMD-4084). The porcine complex I structure is from the respirasome structure of Yang and coworkers (22) (5GUP.pdb and EMD-9539).

**Supplementary Table 1-.**
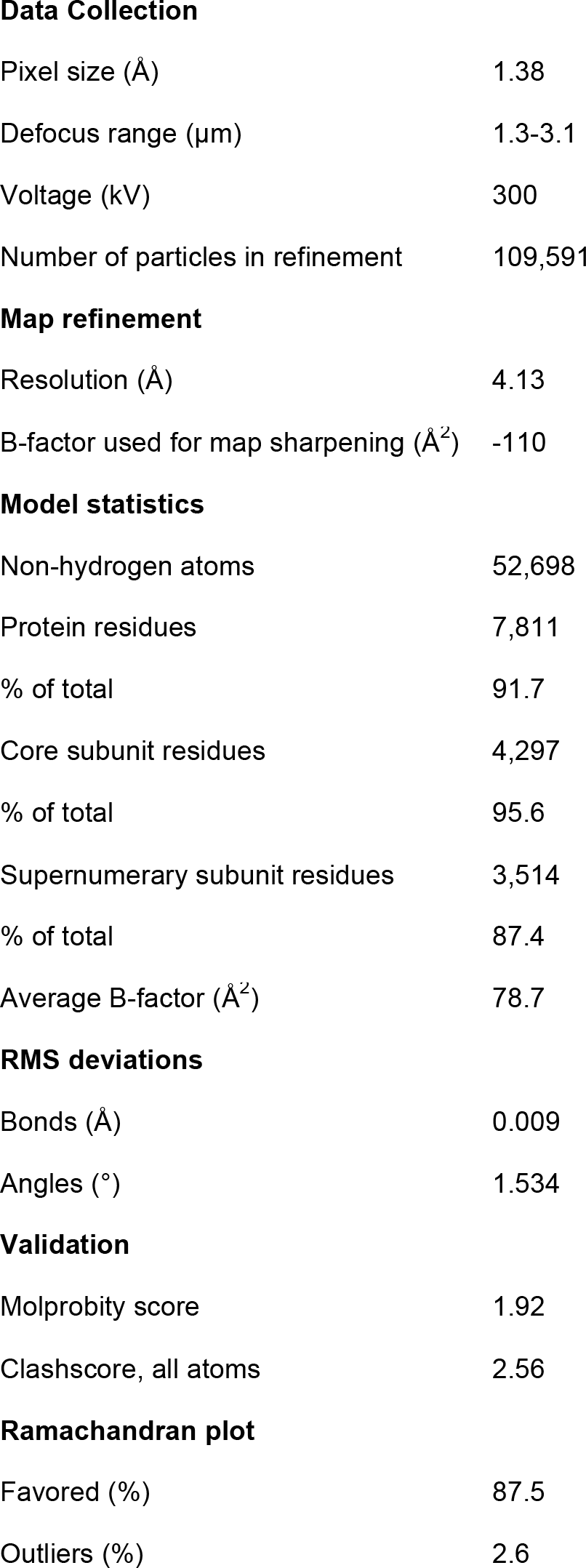
Data collection, refinement, and model statistics for the dominant class of the deactive enzyme.

|  |  |
| --- | --- |
| Pixel size (Å) | 1.38 |
| Defocus range (μm) | 1.3-3.1 |
| Voltage (kV) | 300 |
| Number of particles in refinement | 109,591 |
| **Map refinement** |  |
| Resolution (Å) | 4.13 |
| B-factor used for map sharpening (Å<sup>2</sup>) | -110 |
| **Model statistics** |  |
| Non-hydrogen atoms | 52,698 |
| Protein residues | 7,811 |
| % of total | 91.7 |
| Core subunit residues | 4,297 |
| % of total | 95.6 |
| Supernumerary subunit residues | 3,514 |
| % of total | 87.4 |
| Average B-factor (Å<sup>2</sup>) | 78.7 |
| **RMS deviations** |  |
| Bonds (Å) | 0.009 |
| Angles (°) | 1.534 |
| **Validation** |  |
| Molprobity score | 1.92 |
| Clashscore, all atoms | 2.56 |
| **Ramachandran plot** |  |
| Favored (%) | 87.5 |
| Outliers (%) | 2.6 |

**Supplementary Table 2-.** Summary of the models for the core subunits of deactive bovine complex I.

| Subunit | Other names* | Chain | Total residues | Modelled residues | Assigned residues | Unknown residues | % residues modelled | % residues assigned | % with sidechains | % unknown residues |
| --- | --- | --- | --- | --- | --- | --- | --- | --- | --- | --- |
| ND1 | Nqo8<br>NuoH | H | 318 | 3-200<br>218-315 | 3-200<br>218-315 | - | 93.1 | 93.1 | 90.3 | 0 |
| ND2 | Nqo14<br>NuoN | N | 347 | 2-345 | 2-345 | - | 99.1 | 99.1 | 87.3 | 0 |
| ND3 | Nqo7<br>NuoA | A | 115 | 2-27<br>51-112 | 2-27<br>51-112 | - | 76.5 | 76.5 | 72.2 | 0 |
| ND4 | Nqo13<br>NuoM | M | 459 | 3-459 | 3-459 | - | 99.6 | 99.6 | 93.0 | 0 |
| ND4L | Nqo11<br>NuoK | K | 98 | 2-96 | 2-96 | - | 96.9 | 96.9 | 96.9 | 0 |
| ND5 | Nqo12<br>NuoL | L | 606 | 2-605 | 2-605 | - | 99.7 | 99.7 | 86.1 | 0 |
| ND6 | Nqo10<br>NuoJ | J | 175 | 2-172 | 2-172 | - | 97.7 | 97.7 | 80.0 | 0 |
| 75 kDa | NDUFS1<br>Nqo3<br>NuoG | G | 704 | 6-693 | 6-693 | - | 97.7 | 97.7 | 29.0 | 0 |
| 51 kDa | NDUFV1<br>Nqo1<br>NuoF | F | 444 | 14-438 | 14-438 | - | 95.7 | 95.7 | 18.2 | 0 |
| 49 kDa | NDUFS2<br>Nqo4<br>NuoCD | D | 430 | 5-50<br>61-430 | 5-50<br>61-430 | - | 96.7 | 96.7 | 87.0 | 0 |
| 30 kDa | NDUFS3<br>Nqo5<br>NuoCD | C | 228 | 10-213 | 10-213 | - | 89.5 | 89.5 | 86.0 | 0 |
| 24 kDa | NDUFV2<br>Nqo2<br>NuoE | E | 217 | 8-193 | 8-193 | - | 85.7 | 85.7 | 1.8 | 0 |
| PSST | NDUFS7<br>Nqo6<br>NuoB | B | 179 | 27-173 | 27-173 | - | 82.1 | 82.1 | 82.1 | 0 |
| TYKY | NDUFS8<br>Nqo9<br>NuoI | I | 176 | 1-176 | 1-176 | - | 100 | 100 | 92.0 | 0 |

\*The names of the human, *T. thermophilus* and *E. coli* subunits (if different to the names in *B. taurus*).

**Supplementary Table 3-.** Summary of the models for the supernumerary subunits of deactive bovine complex I.

| Subunit | Human name | Chain | Total residues | Modelled residues | Assigned residues | Unknown residues | % residues modelled | % residues assigned | % with sidechains | % unknown residues |
| --- | --- | --- | --- | --- | --- | --- | --- | --- | --- | --- |
| 42 kDa | NDUFA10 | O | 320 | 5-318 | 5-318 | - | 98.1 | 98.1 | 26.6 | 0 |
| 39 kDa | NDUFA9 | P | 345 | 2-186<br>200-252<br>280-324 | - | 2-186<br>200-252<br>280-324 | 82.0 | 0 | 0 | 82.0 |
| 18 kDa | NDUFS4 | Q | 133 | 11-133 | 11-133 | - | 91.7 | 92.5 | 0 | 0 |
| 15 kDa | NDUFS5 | e | 105 | 6-94 | 6-94 | - | 84.8 | 84.8 | 42.9 | 0 |
| 13 kDa | NDUFS6 | R | 96 | 1-93 | 1-93 | - | 96.9 | 96.9 | 34.3 | 0 |
| 10 kDa | NDUFV3 | s | 75 | 34-74 | 34-74 | - | 54.7 | 54.7 | 0 | 0 |
| AGGG | NDUFB2 | j | 72 | 8-59 | - | 8-59 | 72.2 | 0 | 0 | 72.2 |
| ASHI | NDUFB8 | l | 158 | 5-122 | - | 5-122 | 74.7 | 0 | 0 | 74.7 |
| ESSS | NDUFB11 | g | 125 | 25-121 | 25-121 | - | 77.6 | 77.6 | 41.6 | 0 |
| KFYI | NDUFC1 | c | 49 | 1-46 | 1-46 | - | 93.9 | 93.9 | 59.2 | 0 |
| MNLL | NDUFB1 | f | 57 | 3-56 | 3-56 | - | 94.7 | 94.7 | 42.1 | 0 |
| MWFE | NDUFA1 | a | 70 | 1-64 | 1-64 | - | 91.4 | 91.4 | 71.4 | 0 |
| PDSW | NDUFB10 | p | 175 | 4-172 | 76-142 | 4-75<br>143-172 | 96.6 | 38.3 | 30.9 | 58.3 |
| PGIV | NDUFA8 | X | 171 | 5-168 | 5-168 | - | 95.9 | 95.9 | 53.8 | 0 |
| SDAP-α | NDUFAB1 | T | 88 | 8-82 | 8-82 | - | 85.2 | 85.2 | 0 | 0 |
| SDAP-β | NDUFAB1 | U | 88 | 4-88 | 4-88 | - | 96.6 | 96.6 | 0 | 0 |
| SGDH | NDUFB5 | h | 143 | 7-140 | 7-140 | - | 93.7 | 93.7 | 21.7 | 0 |
| B22 | NDUFB9 | n | 178 | 10-175 | 10-175 | - | 93.3 | 93.3 | 36.0 | 0 |
| B18 | NDUFB7 | o | 136 | 57-114 | 57-114 | - | 42.6 | 42.6 | 2.9 | 0 |
| B17.2 | NDUFA12 | q | 145 | 2-139 | 2-139 | - | 95.2 | 95.2 | 0 | 0 |
| B17 | NDUFB6 | i | 127 | 6-32<br>40-118 | 6-32 | 40-118 | 83.5 | 21.3 | 14.2 | 62.2 |
| B16.6 | NDUFA13 | Z | 143 | 5-141 | 5-98 | 99-141 | 95.8 | 65.7 | 47.6 | 30.1 |
| B15 | NDUFB4 | m | 128 | 11-128 | 11-128 | - | 92.2 | 92.2 | 67.2 | 0 |
| B14.7 | NDUFA11 | Y | 140 | 1-138 | 1-138 | - | 98.6 | 98.6 | 98.6 | 0 |
| B14.5a | NDUFA7 | r | 112 | 1-69<br>93-111 | - | - | 78.6 | 78.6 | 0 | 0 |
| B14.5b | NDUFC2 | d | 120 | 4-116 | 4-97 | 98-116 | 94.2 | 78.3 | 52.5 | 15.8 |
| B14 | NDUFA6 | W | 127 | 16-126 | 16-126 | - | 87.4 | 87.4 | 57.5 | 0 |
| B13 | NDUFA5 | V | 115 | 8-113 | 8-113 | - | 92.2 | 92.2 | 40.9 | 0 |
| B12 | NDUFB3 | k | 97 | 16-89 | - | 16-89 | 76.3 | 0 | 0 | 76.3 |
| B9 | NDUFA3 | b | 83 | 1-80 | 1-45 | 46-80 | 96.4 | 54.2 | 54.2 | 42.2 |
| B8 | NDUFA2 | S | 98 | 16-95 | 16-95 | - | 81.6 | 81.6 | 0 | 0 |

